# Structurally distinct polymorphs of Tau aggregates revealed by nanoscale infrared spectroscopy

**DOI:** 10.1101/2021.08.12.456130

**Authors:** Siddhartha Banerjee, Ayanjeet Ghosh

## Abstract

Aggregation of the tau protein plays a central role in several neurodegenerative diseases collectively known as tauopathies, including Alzheimer’s and Parkinson’s disease. Tau misfolds into fibrillar beta sheet structures that constitute the paired helical filaments found in Neurofibrillary tangles. It is known that there can be significant structural heterogeneities in tau aggregates associated with different diseases. However, while structures of mature fibrils have been studied, the structural distributions in early stage tau aggregates is not well understood. In the present study, we use AFM-IR to investigate nanoscale spectra of individual tau fibrils at different stages of aggregation and demonstrate the presence of multiple fibrillar polymorphs that exhibit different secondary structures. We further show that mature fibrils contain significant amounts of antiparallel beta sheets. Our results are the very first application of nanoscale infrared spectroscopy to tau aggregates and underscore the promise of spatially resolved infrared spectroscopy for investigating protein aggregation.

## Introduction

The misfolding and aggregation of tau proteins into fibrillar aggregates is the pathological hallmark of many neurodegenerative diseases called tauopathies, including Alzheimer’s and Parkinson’s disease^1–5^. Tau is a microtubule (MT)-associated protein that misfolds into insoluble cellular deposits called Neurofibrillary Tangles (NFTs)^1, 2, 5, 6^. While early evidence pointed to the potential neurotoxicity of NFTs, it is now believed that prefibrillar oligomeric assemblies are the main neurotoxic species^3, 5, 6^. Elucidation of specific tau fibrillization pathways may provide insights into disease mechanisms and reveal potential therapeutic targets for drug discovery. Significant efforts have been invested to understand the aggregation of tau and the role of different factors that modulate the aggregation^1, 2, 7–15^. It is known that tau filaments in NFTs have a cross beta structure similar to amyloid plaques. However, comprehensively elucidating the structural evolution of tau that results in fibril formation remains a challenge. In the human brain, six different tau isoforms have been identified that differ with respect to the number of amino acid resides^2, 5^. The dominant isoform and fibrillar structure can vary with disease^2^. All the tau isoforms are large polypeptides, and therefore exhibit structural flexibility. Furthermore, tau and its isoforms exhibit polymorphism: essentially the same peptide aggregates into different fibrillar structures, that differ not only with respect to morphology but also with respect to molecular arrangement^8, 9, 16^. All of the above factors make isolation and structural analysis of specific tau aggregates formed at different stages of the aggregation a difficult task. The nature of amyloid aggregation in general is such that it generates a plethora of aggregated species which are transient in nature and are in equilibrium with each other. Recently cryo-EM has been successfully applied for solving tau fibrillar structures; however its applications remain limited to essentially mature fibrils that are the end point of aggregation, and not intermediates^9, 10^. In particular, the structural aspects of early-stage tau intermediates are not well understood. The gold-standard for determining secondary structure of amyloid-like aggregates in general are spectroscopic techniques like solid state Nuclear Magnetic Resonance (ssNMR) and Fourier Transform Infrared Spectroscopy (FTIR)^15, 17–19^. However, none of these techniques are capable of providing spatial resolution on the scale of individual aggregates, and without spatial resolution, it is difficult to unequivocally attribute spectral features to specific aggregates or morphologies or determine which transient species evolve into a particular morphology. Essentially it is not possible to determine if observed spectral features arise from a particular aggregation state or a statistical mixture of different conformations.

As a result, only average structures of oligomers and fibrils are known, and structural variations and heterogeneities within each state, if any, are not well understood. To advance our knowledge of aggregation of tau and other amyloidogenic proteins under different conditions, it is imperative to understand the structure of each member of the conformational ensemble at different stage of aggregation. The past decade has seen the development of novel approaches for improving the spatial resolution of vibrational spectroscopy that combine infrared spectroscopy to AFM to achieve nanometer scale resolution. One of the most recently developed AFM-based approaches takes advantage of photothermal-induced resonance (PTIR), where the local thermal expansion resulting from infrared absorption by a sample is sensed by an AFM tip^20–23^. Photothermal AFM-IR thus bypasses resolution limits in conventional IR microscopy by using the tip of an AFM probe to measure infrared absorption. Absorption of radiation through resonant excitation of an infrared mode leads to a thermal expansion of the sample, producing an impulsive force on the AFM cantilever. The resultant AFM probe response is proportional to the infrared absorbance, and scanning the wavelength yields an infrared absorption spectrum that corresponds to a nanoscale region of the sample (Fig. 1). Thus, unlike conventional optical techniques, AFM-IR can examine nanoscale structures at unprecedented chemical detail and combines the best of both worlds: the spatial resolution of AFM and the chemical resolution of infrared. While AFM-IR has been employed to study aggregates of amyloidogenic peptides like beta amyloid and alpha synuclein^23–26^, it has never been applied to study aggregation of tau. In this study, we leverage the capabilities of AFM-IR and investigate fibrils of the tau-441 isomer at different stages of aggregation to show that at early stages of aggregation, there is significant heterogeneity in fibrillar structure even without any significant variations in morphology. Our results demonstrate for the very first time that there can be underlying differences in secondary structure of fibrils that exhibit the same morphology. Multiple structurally distinct fibril polymorphs are observed; one more structurally ordered than the others, which is also transient in nature and is not found in mature fibrils. Furthermore, we also demonstrate that tau fibrils can contain antiparallel beta sheets, which is typically not associated with fibrillar morphologies.

**Figure 1:**
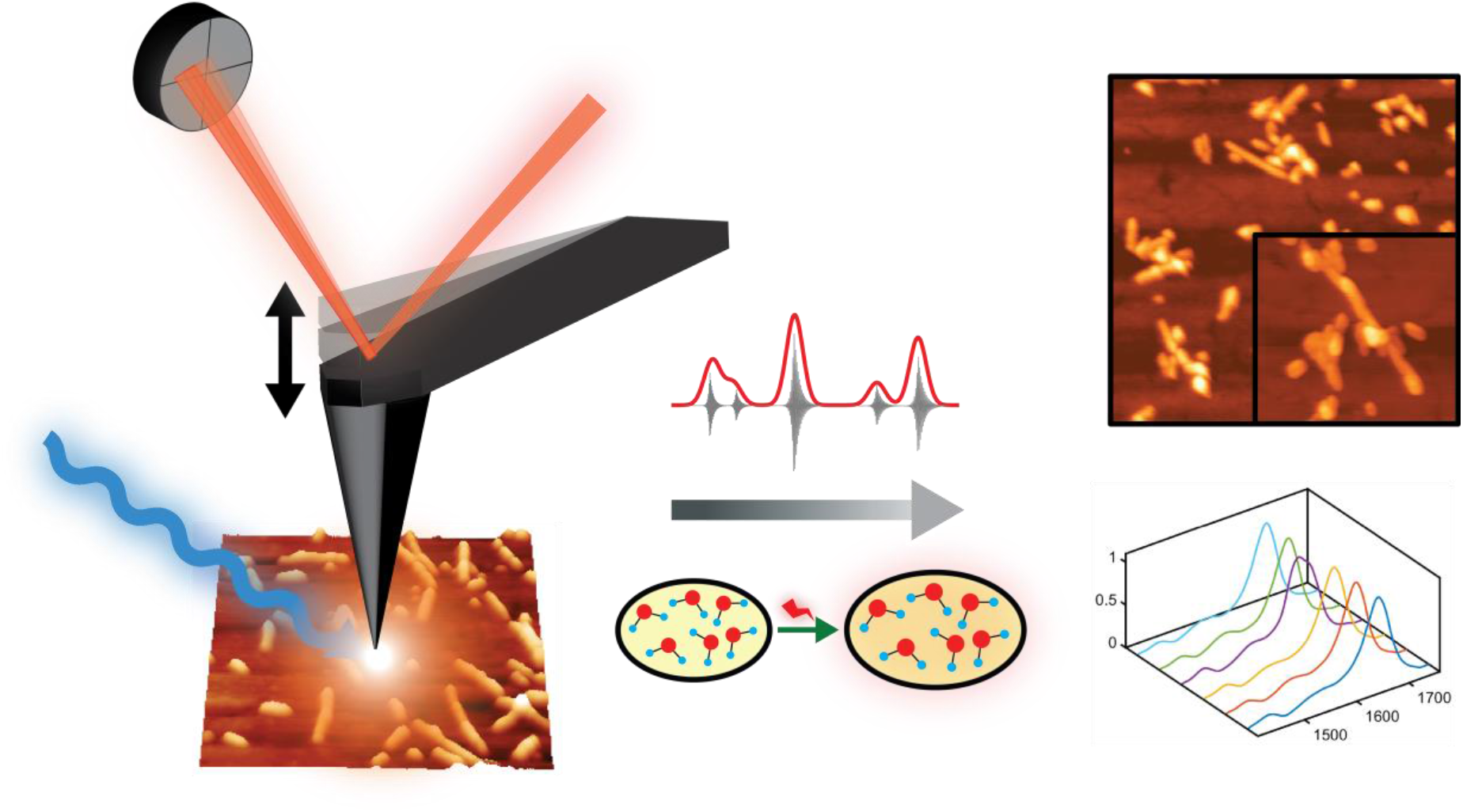
Schematic representation of AFM-IR experiment. Nanoscale IR measurements are performed on the same tau fibril which are visualized by AFM imaging, providing both morphological and structural information. Thermal expansions arising from resonant infrared excitation is sensed by the AFM cantilever, generating nanoscale IR spectra.

## Results and discussion

To understand the structural evolution of fibrils with time, we investigated fibrils at different time points of aggregation, namely after 3, 5, 10 and 15 days of aggregation. For each measurement, aliquots were deposited on gold substrates and dried under nitrogen. AFM topographic images of tau fibrils after 3 days of incubation at 37°C are shown in Fig. 2. Fig. 2A shows a representative fibril cluster. More AFM images of individual fibrils at all the different aggregation stages is shown in the Supporting Fig. 1-5. To obtain the overall height distribution of the fibrils in the sample, height values of the individual fibrils were measured and plotted as a histogram (Fig. 2B). The data was fitted with a Gaussian to determine the mean height value of 6.3±0.7 nm. AFM images of 5-day fibrils and corresponding height analysis is shown in Figure 2C-D. Individual fibrils along with fibril clusters are observed. The height of the fibrils is 6.5±0.9 nm which remains close to that of 3-day fibril sample. Next, tau fibrils with 10-day incubation were probed (Fig. 2E-F). Individual fibrils were observed on the gold surface (Fig. 2E). The average height of 10-day fibrils have been found to be 7.2±1.0 nm. Mature tau fibrils, generated after 15 days of aggregation, are shown in Fig. 2G. The 15-day fibrils are long and tangled with each other generating a network like morphology. At this stage of aggregation, separate fibrils are no longer visible. The average fibril height is 42.5±40.5 nm (Figure 2H) which is significantly higher compared to the other fibrils those has been generated at the earlier time points. The broad distribution of fibrils height indicates that mature fibrils have varied height values, as opposed to the narrow distribution of earlier fibrils. The fibrillar structures observed here are consistent with previous AFM reports of tau aggregation^11, 27, 28^. Another important observation from the AFM measurements is that the tau fibrils generated in solution are homogeneous in their morphology. They are straight, lacking any kind of branching or twisted conformations, with minimal height changes within one fibril. With fibril maturation, we see an increase in fibril height, but no new morphologies develop. This can be clearly visualized in Supplementary Fig. 5, where 3-day and 10-day fibril morphologies are shown in a rainbow color map, demonstrates that they have minimal height changes within one fibril.

**Figure 2:**
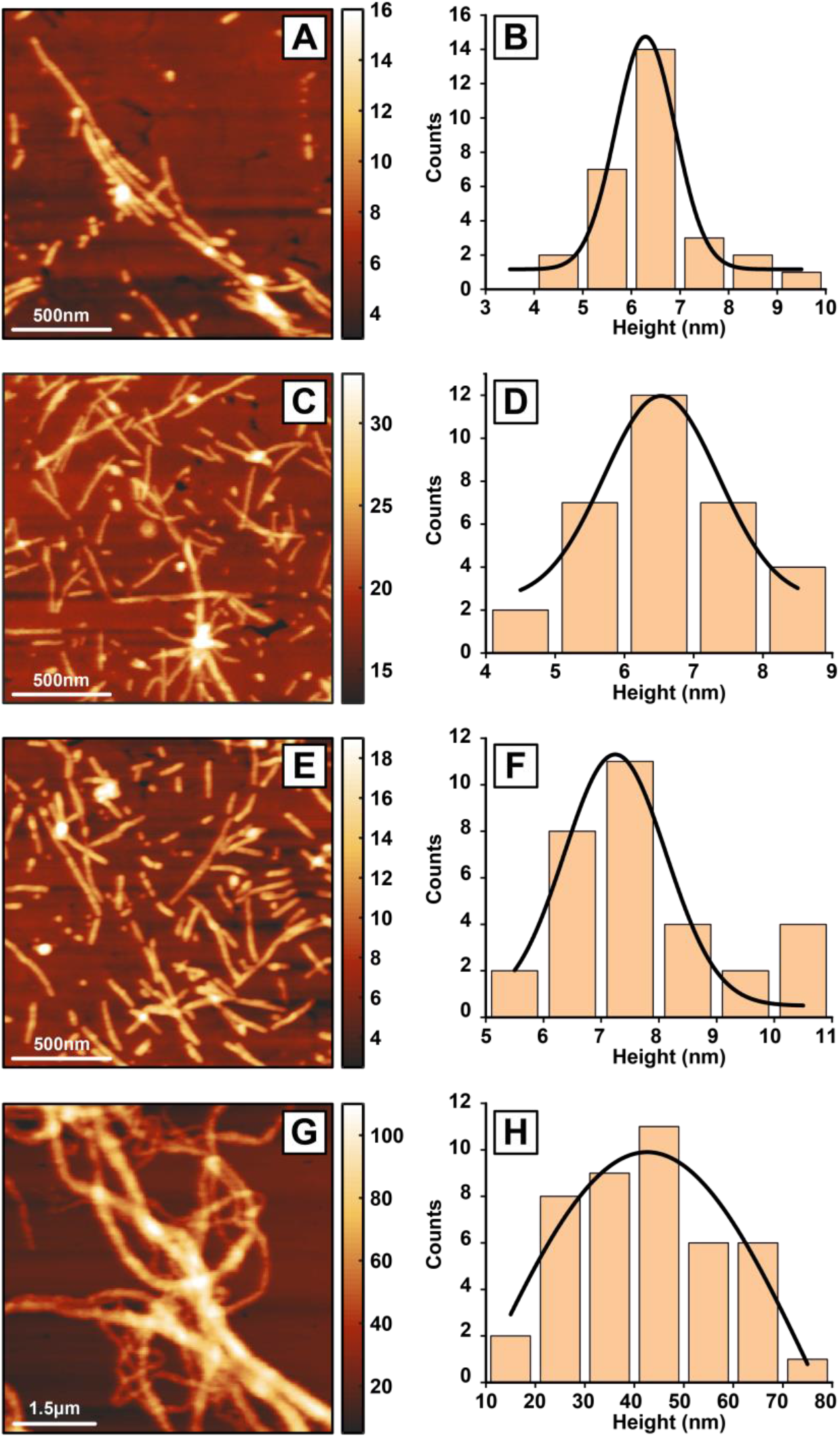
AFM imaging of tau fibrils. Tau fibrils have been characterized at different time periods of 3 days (A-B), 5 days (C-D), 10 days (E-F) and 15 days (G-H). AFM topographic images of the fibrils for 3 days (A), 5 days (C), 10 days (E) and 15 days (G) show the fibril morphologies. The height values, as indicated by the colorbars, are in nanometers. The height value of the fibrils was measured from several fibrils from each time periods of 3 days (B), 5 days (D), 10 days (F), 15 days (H) and plotted as a histogram. The distributions are fitted with a Gaussian to determine the peak height value in each case. The fibril height for 3 days, 5 days, 10 days and 15 days are 6.3±0.7 nm, 6.5±0.9 nm, 7.2±1.0 nm and 42.5±40.5 nm, respectively.

While the fibrils do not exhibit any significant morphological differences, the nanoscale IR spectra of fibrils contain significant differences. Specifically, for the 3-day fibrils we observe three different fibril polymorphs with distinct spectra. For 5-day, 10-day and 15-day matured fibrils, we observe only one conformation. We note that typically polymorphism is used to signify morphologically distinct fibrils that can also be different with respect to molecular structure^2, 16^; in our case the morphology is invariant, but the fibrils can be categorized into three distinct subtypes based on their spectra. For clarity, we refer to these structures/polymorphs as Type 1, Type 2 and Type 3 respectively for the rest of this article. Representative infrared spectra in the amide-I range are shown in Fig. 3. The spectra are the average of multiple measurements taken along the length of the fibrils. More spectra are provided in the Supplementary Information (Fig. S6). The spectrum of 3-day fibril type-1 contains a prominent peak at ~1628cm^−1^ and a smaller shoulder at ~1670cm^−1^. Additionally, a weak band can be seen at ~1740cm^−1^. The spectrum of fibril type-2 is significantly different and contains a broadened asymmetric amide-I band centered at ~1650cm^−1^. No significant intensity is observed above 1700cm^−1^. For fibril type-3, we observe an amide-I spectrum that is very similar to type-2, with one major exception: presence of a prominent band at ~1738cm^−1^. It is interesting to note that the fibril spectra, with the exception of the type-1 polymorph, do not contain a sharp peak at typical beta sheet frequencies of ~1630cm-1, even though fibrillar aggregates are known to be in-register parallel beta sheets. The broader linewidth observed for type 2 and type 3 fibrils, in addition to the lack of a sharp peak at beta sheet frequencies would indicate presence of structural disorder. Interestingly, we did not find any significant variance in spectra along a single fibril (Supplementary Fig. 5), indicating that the fibrils are well defined structures, and have intrinsic structural disorder.

**Figure 3:**
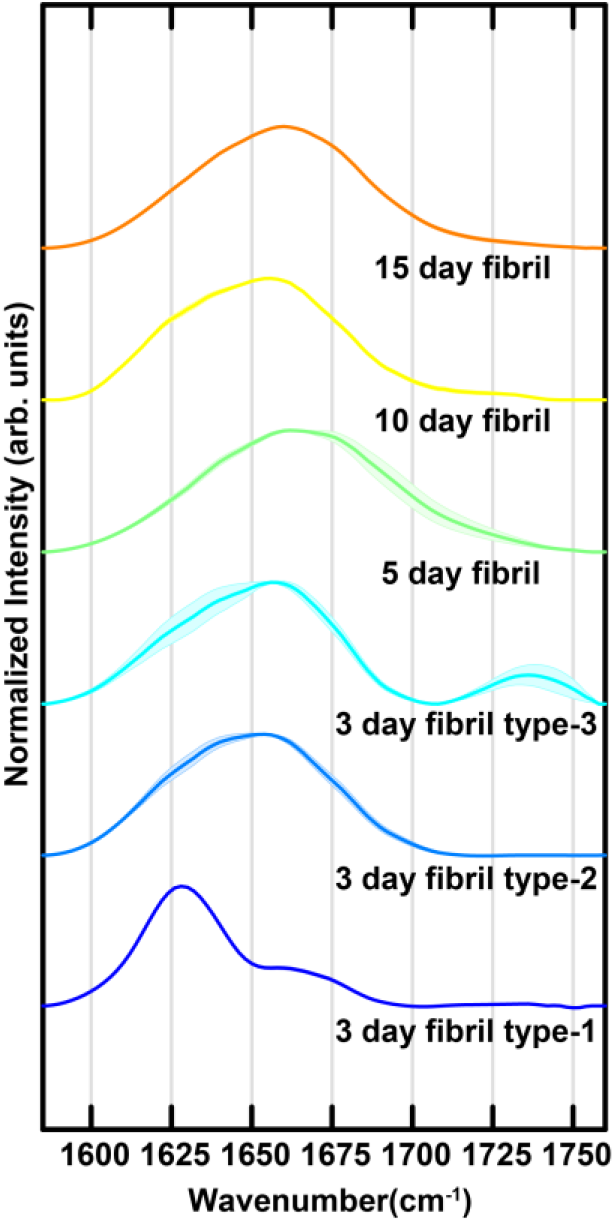
Infrared spectra of tau fibrils. Mean IR spectra of tau fibrils were measured at along the length of different fibrils at time periods of 3 days, 5 days, 10 days and 15 days. Different 3-day fibrils exhibit distinctly unique spectra, labeled as type-1, type-2 and type-3. The shaded areas represent the standard deviation of spectral intensity.

The 5-day, 10-day and 15-day mature fibrils do not exhibit any spectral heterogeneity, and a single polymorph is observed for each case (Fig. 3). The 5-day fibril spectrum is slightly shifted (~6cm^−1^) to higher wavenumbers compared to the type 2 and type 3 fibrils and also exhibits increased linewidth. The 10-day and 15-day matured fibrils have spectra are similar to that of the type 2 fibrils, with the latter shifted to higher wavenumbers by ~4cm^−1^. All of the 5-day, 10-day and 15-day fibrils lack a distinct peak above 1700cm^−1^.

Although the spectra above clearly indicate evolution of the fibrils, it is difficult to precisely understand the exact underlying structural changes from inspection of the spectra alone. Therefore, to gain further insights into the spectral and structural variations between the polymorphs observed, the spectra were deconvoluted through spectral fitting. Amide I spectra of proteins usually contain contributions from different secondary structures^29, 30^. Since the secondary structure contributing to each of the observed spectra is not exactly known a priori, we turned to the second derivative spectra to determine the number of peaks (Supplementary Fig. 7). Using the second derivative of spectral data to determine underlying peaks is a well-known practice in spectroscopy^31, 32^. We used the number of peaks in the second derivative spectra and their corresponding frequencies as a starting point for spectral fitting. The fit results for the mean spectra of the 3-day, 5 day, 10-day and 15-day mature fibrils is shown in Fig. 4. The fit parameters are provided as Supplementary Table 1. For the type-1 3-day fibrils, three Gaussian bands fit both spectra with high fidelity, with center frequencies at 1628cm^−1^, 1659cm^−1^ and 1670cm^−1^. The type-2 3-day fibril spectrum fits to five underlying bands, with frequencies 1626cm^−1^, 1642cm^−1^, 1662cm^−1^, 1680cm^−1^ and 1694cm^−1^. As expected from the spectral similarity, the type-3 fibril fits to the same 5 bands but also requires an additional peak at 1736cm^−1^. Extending our spectral deconvolution analysis to more mature fibrils, we see that 5-day, 10-day and 15-day fibrils all essentially contain the same distribution of secondary structures. The 5-day, 10-day and 15-day fibril spectra all require six bands for optimal fitting, wherein the high wavenumber peak shifts to ~1725cm^−1^ and the other 5 peaks are similar to the type-2 fibrils.

**Figure 4:**
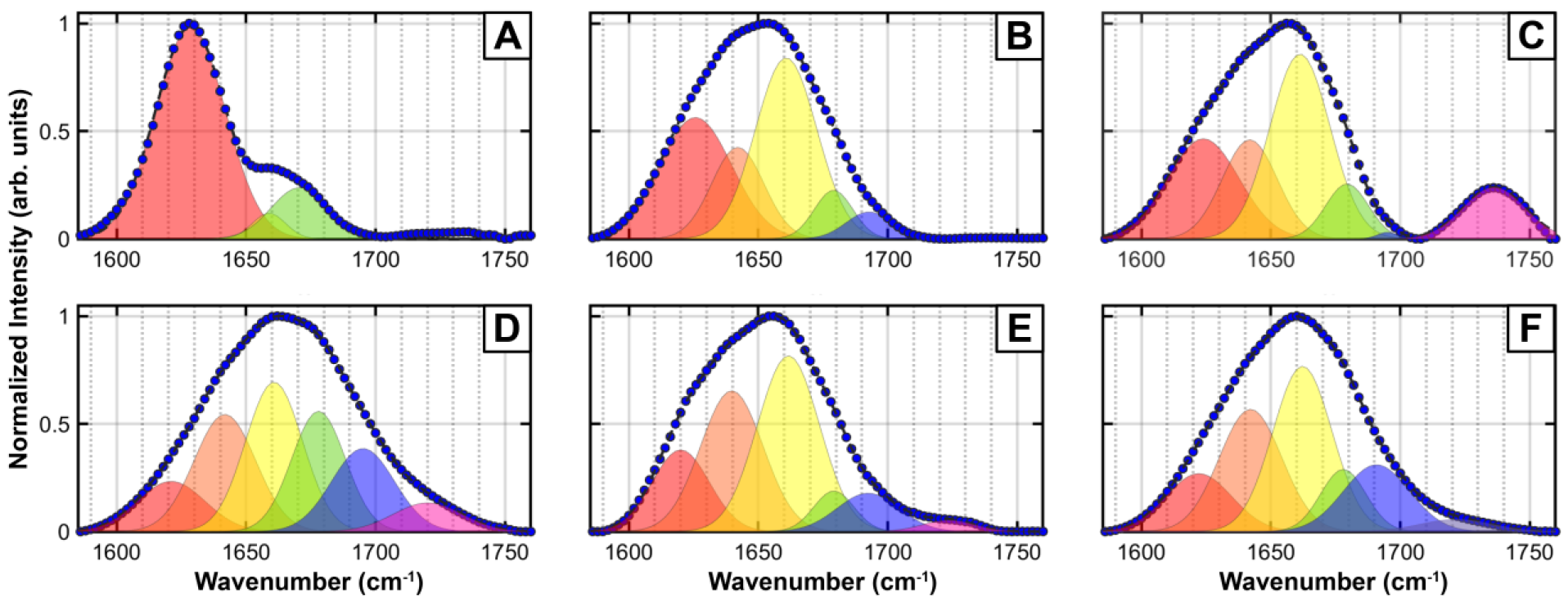
Deconvolution of fibril spectra. Spectral fitting was performed for the different spectrally unique types of tau fibrils identified at different aggregation times. A. 3-day fibril type-1, B. 3-day fibril type-2, C. 3-day fibril type-3, D. 5 day fibril, E. 10-day fibril and F. 15-day fibril. The fit parameters are provided in the Supporting Information. The measured spectra are shown as filled circles, while the dashed lines represent the fits. The individual bands contributing to the overall spectra are shown as shaded peaks.

The 1626-1628cm^−1^ peak in all of the fits can be attributed to beta sheets, indicating that all fibrillar polymorphs contain beta sheet structure^17, 19, 29, 30^. The1642cm^−1^ peak typically arises from random coils, while the peaks at ~1660cm^−1^ and 1682cm^−1^ are typically assigned to beta turns^29, 30^. Taken together, the spectral deconvolution indicates that upon maturation, there is more disorder in the fibril structure, as evidenced by the increase in the random coil peak relative to the beta sheet peak. The presence of beta turns and their relative increase, evidenced by the intensity of the 1660cm^−1^ fitted peak, is consistent with the expected cross beta structure of the fibrils. Presence of a bands at ~1694cm^−1^ and ~1725cm^−1^, however, are somewhat unexpected. The former has been typically attributed to antiparallel beta sheets^29, 30, 33^ and has been observed in beta amyloid^33, 34^ and alpha synuclein oligomers^26^. While antiparallel beta sheet structures are known for oligomeric amyloid assemblies, their existence in fibrils has been rarely observed. 2D IR studies have identified antiparallel beta sheet signatures in synuclein fibrils^35^; specific mutants of beta amyloid also exhibit antiparallel beta sheet structure^35^. Our results, to the best of our knowledge, are the very observation of antiparallel beta sheets for tau aggregates. The antiparallel beta sheet peak is conspicuously absent in the type-1 polymorph. This suggests that fibrils can adopt either a rigid well-ordered structure (polymorph type-1) that is primarily parallel beta sheets but increased structural flexibility and/or disorder can lead to formation of antiparallel beta sheets. However, it is important to note that the ~1626cm^−1^ peak does not typically shift significantly between parallel and antiparallel beta sheets. Our results therefore do not preclude the possibility of parallel beta sheets in any of the fibrils where the ~1694cm^−1^ peak is observed. The peak at ~1725cm^−1^ cannot be attributed to a backbone amide vibration, and most likely arises from COOH stretch of sidechain carboxylic acids^29, 30^. The Tau 441 peptide sequence has multiple carboxylic acids, including in the microtubule binding repeat domains, which have been shown to have high propensity for forming beta sheets^5, 6^. In fact, AFM-IR measurements of other amyloid aggregates have identified similar peaks^24^, which have been attributed to carboxylic acids. However, this carboxylic acid band is more prominent in some fibrils than others. In infrared spectroscopy, dipolar alignment in ordered structures often leads to increased intensity of absorption bands; a relevant example is the formation of ordered beta sheets from unordered peptide, leading to sharp intense bands at ~1625cm^−1^. Thus, the presence of intense carboxylic acid bands possibly arises from structural ordering of glutamic and aspartic acid sidechains. However, it is important to note in this context that AFM-IR measurements are different from isotropic infrared spectra acquired in solution, and the spectra measured in AFM-IR is a convolution of laser polarization and illumination configuration^36, 37^ Therefore, it is difficult to precisely identify the structural underpinnings of this peak, and more research is necessary to determine their exact origins. We aim to address this in future work.

The aggregation of tau isomers *in vitro* has been investigated in detail; however, the structure of early and/or transient intermediates is not very well known. Oligomeric species in amyloid protein aggregation have been identified to be either on or off pathway towards fibril formation. Since fibrils are believed to be the endpoint of aggregation, fibril structures, including different polymorphs thereof, are not generally thought of as transient or off pathway. Heterogeneity in fibrillar structure is known to exist but has been known to present itself concurrently with morphological variations^2, 8, 16, 38^. Our results are unique in that they point to variations in secondary structure of fibrils even when there are no perceptible morphological differences. We note that with the exception of fibril type-1, all the fibrillar spectra contain the same set of underlying bands, indicating that the type-1 fibril is a transient intermediate that eventually undergoes structural reorganization with maturation. Another possibility is that these ordered parallel beta sheet polymorphs represent an ‘off-pathway’ structure, it must disintegrate into monomeric or prefibrillar aggregates to be reintegrated into ‘on-pathway’ fibrils. We did not find any significant presence of non-fibrillar deposits in the specimens studied; however, the ones present did have a spectrum that resemble the disordered fibrils more (Supplementary Fig. 8). This is in agreement with the off-pathway hypothesis and would thus suggest that the other 3-day polymorphs observed can be viewed upon as on pathway aggregates. However, more detailed analysis of aggregation kinetics is necessary to confirm this hypothesis. The other intriguing observation enabled by AFM-IR is the identification of antiparallel beta sheets in matured fibrils. It is generally believed, and has been evidenced by many studies, that early-stage amyloid aggregates can contain antiparallel structure, which gets converted to parallel beta sheet secondary structure in mature aggregates. We observe an opposite trend: early-stage aggregates can contain ordered parallel beta sheets while more mature fibrils contain antiparallel bet sheet structure.

In summary, we have demonstrated using nanoscale AFM-IR spectroscopy that tau fibrils can have significant structural variations, particularly in early stages in aggregation. The heterogeneity is manifested in the form of structurally distinct polymorphs, which exhibit similar morphology but different secondary structure. Specifically, we identify a transient ordered parallel beta sheet structure in early-stage fibrils, which upon maturation evolves into a more disordered fibril structure that contains antiparallel beta sheets. These results underscore the need to couple spectroscopy with a spatially resolved technique like AFM, as it is not possible to unequivocally make this determination from spatially averaged techniques like FTIR. The experimental results described in this study shows that the tau fibrillar aggregates are heterogeneous and future work will address whether these polymorphs persist under different aggregation conditions and in seeded aggregation from brain lysates to understand their significance in the context of different tauopathies.

## Acknowledgement

This work was supported by the National Institutes of Health (Award 1 R35 GM138162 to A.G.).

## Methods

### Preparation of Tau fibrils

1 mg/ml Tau-441 (rPeptide, USA) in the buffer containing 50mM MES, pH 6.8, 100 mM NaCl and 0.5 mM EDTA was incubated at 37°C with constant shaking at 1000 rpm on a shaking dry bath (Thermo Scientific). Aliquots were taken out at specific time intervals for preparation of AFM samples. Aggregation was continued until the mature fibril was formed.

### Preparation of samples for AFM-IR experiment

Samples for AFM-IR experiments were prepared on template stripped ultra-flat gold surface (Platypus Technologies, USA). 10 μl of the aggregation mixture was taken out at different time intervals and deposited onto gold surface and incubated for 2 min. The substrate was consequently rinsed with 500 μl milli-q water and dried under a gentle stream of nitrogen gas.

### AFM-Nano IR experiment

AFM imaging was carried out in tapping mode at room temperature with a Bruker NanoIR3 instrument, equipped with a mid-IR Quantum Cascade Laser (MIRcat, Daylight solutions). The nominal resonance frequency and the spring constant of the cantilever are 75±15 kHz and 1-7 N/m, respectively. The scan speed was kept between 0.5-1 Hz. After recording the AFM image, IR spectra were collected at several points along the length of different fibrils. The spectra were recorded with a spectral resolution of 2 cm^−1^. For each spectral point, 128 coadditions were performed, and for each spectrum, 16 co-averages were performed. During all measurements, the instrument was purged with dry air to maintain working relative humidity of <5%.

### Spectral analysis and fitting

All spectral analysis was performed using the MATLAB software package. Spectra were fitted with Gaussian peaks using the curve fitting toolbox in MATLAB. Raw spectra were smoothed using a (3,11) Savitzky-Golay filter and a baseline subtraction was performed prior to spectral fitting. Details of the fit parameters are given in the Supplementary Information.

