## Supplementary Information for "Structurally distinct polymorphs of Tau aggregates revealed by nanoscale infrared spectroscopy"

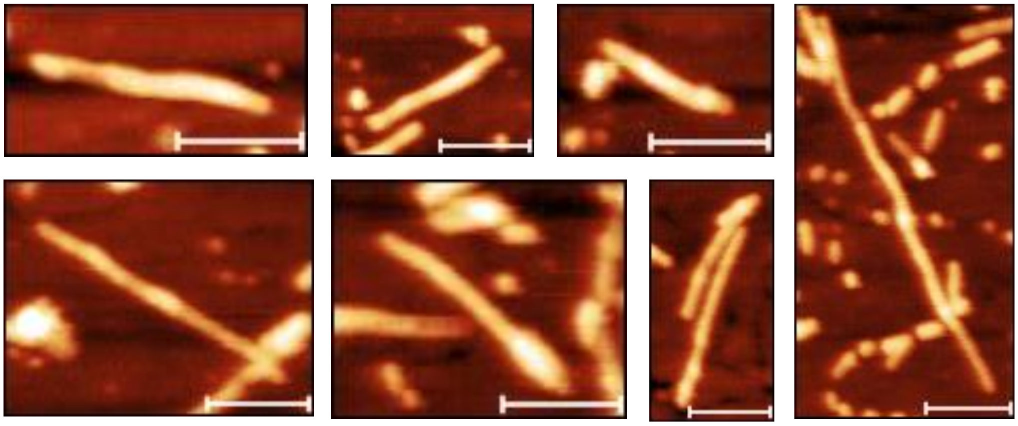

**Supplementary Figure 1: Tau fibril morphology after 3 days**. A gallery of tau fibrils generated after incubating tau monomers for 3 days at 37ºC with shaking. Fibril morphology is homogeneous. No distinct morphology differences have been observed among the fibrils. Scale bar: 200 nm

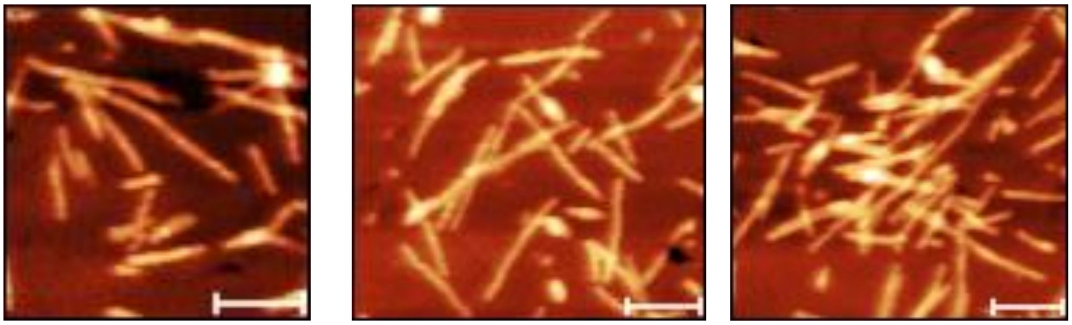

**Supplementary Figure 2: Tau fibril morphology after 5 days**. A gallery of tau fibrils generated while continuing the aggregation for 5 days at 37ºC with shaking. Individual fibrils are visible on the gold substrate. Fibril clusters are observed in some areas. Scale bar: 200 nm

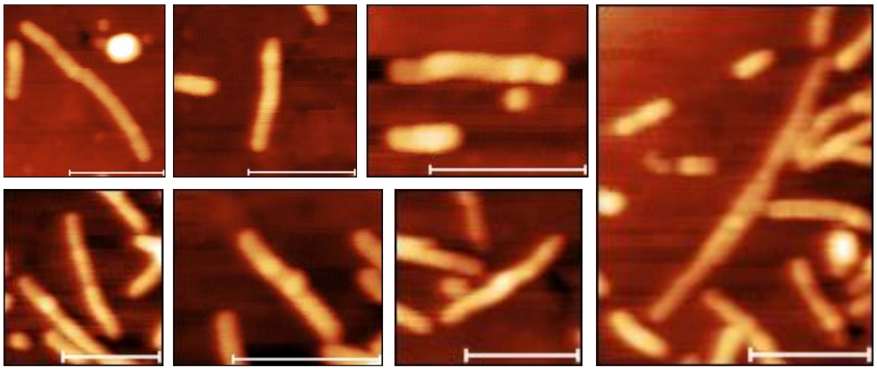

**Supplementary Figure 3: Tau fibril morphology after 10 days**. A gallery of tau fibrils showing the fibril morphology. These fibrils are generated after incubating tau monomers for 10 days at 37ºC with shaking. Single isolated fibrils are observed on the gold substrate. Scale bar: 200 nm

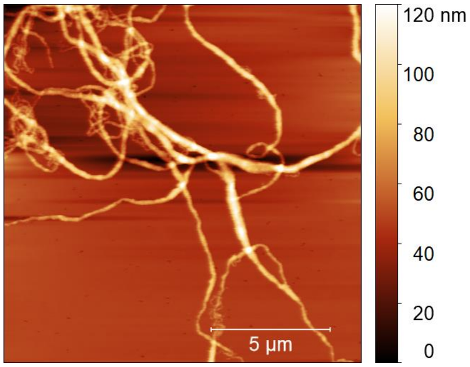


**Supplementary Figure 4: Tau fibril morphology after 15 days**. Large area scan showing the topography of mature tau fibril. Long fibrils with network like morphology is evident.

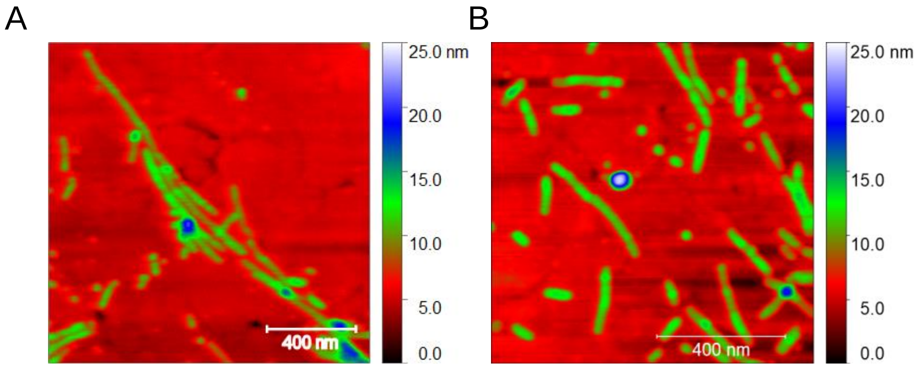


**Supplementary Figure 5: Homogeneous morphology of tau fibrils.** AFM topographic image of tau fibrils with rainbow Z scale after (A) 3 days and (B) 10 days incubation. Same color profile of all the fibrils show that they have very similar height values, although the fibril lengths are different. Fibrils did not show any branching or twisting features.

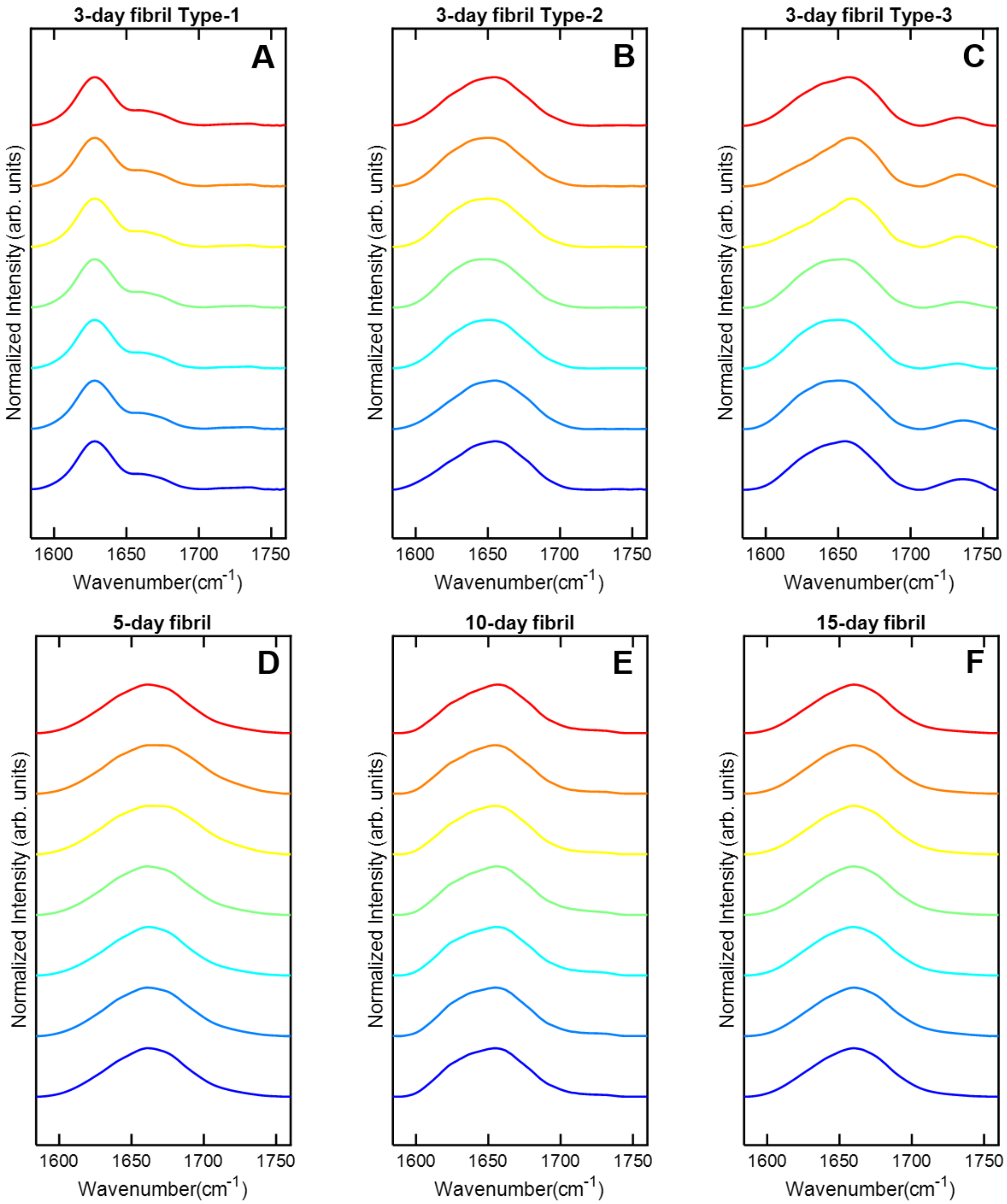


**Supplementary Figure 6: Spectral variations along single tau fibrils.** Representative spectra along single isolated fibrils at different stages incubation. All spectra have been normalized for clarity. While spectral differences can be seen between different fibrils, spectral variations along a single fibril are not significant.

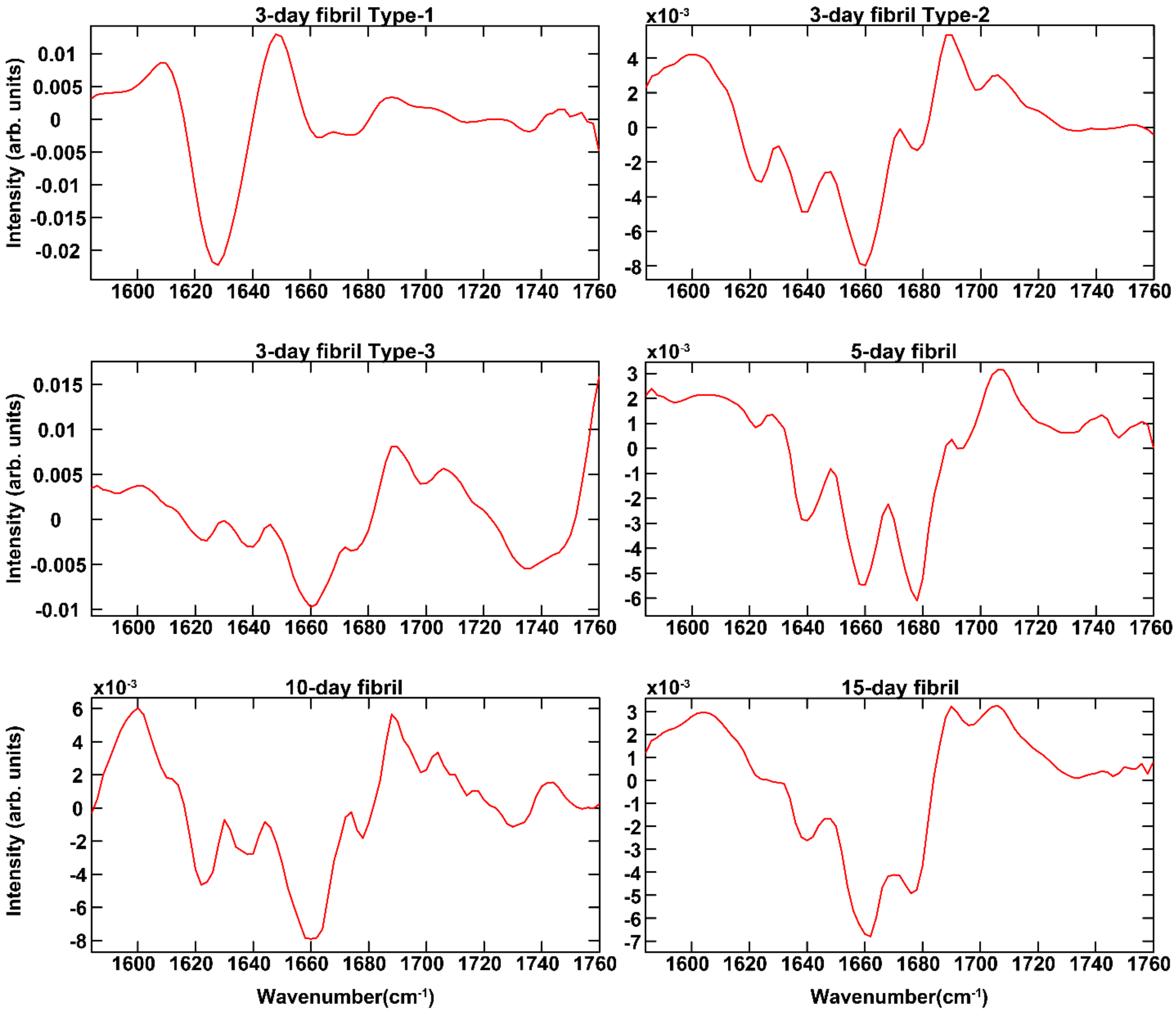


**Supplementary Figure 7: second derivative spectra tau fibrils.** Second derivative of mean spectra of fibrils at different aggregation stages. The derivative spectra were smoothed with a 5-point moving average filter.

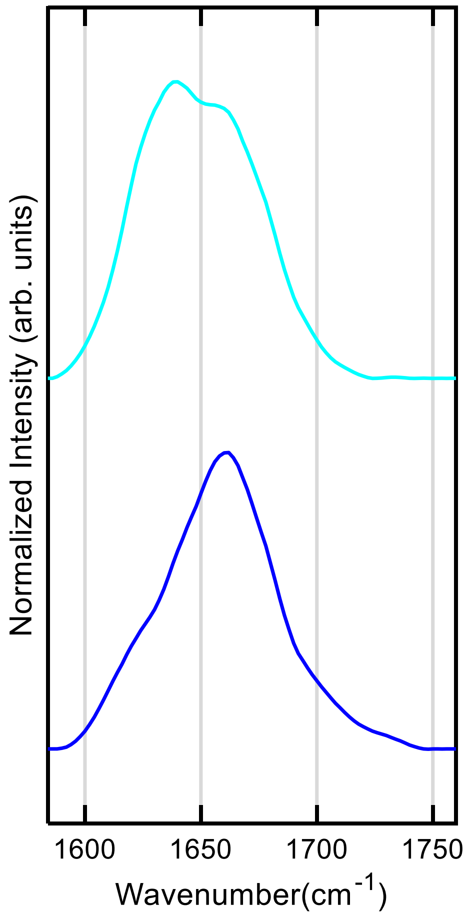


**Supplementary Figure 8: oligomer spectra.** IR spectra of two individual oligomers identified in the AFM scans of 3-day fibril samples.

**Supplementary Table 1. Spectral Fitting parameters**

All spectra were fitted with a sum of Gaussian peaks:

$$S = \sum_i a_i e^{-\left(\frac{x - x_0}{\sigma}\right)^2}$$

Spectra were normalized to the maximum intensity of the amide-I band prior to fitting.

| 3-day fibril Type-1 | Center frequency ($x_0$, $cm^{-1}$) | Amplitude (a) | Width ($\sigma$, $cm^{-1}$) |
|---|---|---|---|
| | 1628.4 | 0.99 | 19.0 |
| | 1658.7 | 0.12 | 9.2 |
| | 1670.4 | 0.24 | 15.6 |
| | - | - | - |

| 3-day fibril Type-2 | Center frequency ($x_0$, $cm^{-1}$) | Amplitude (a) | Width ($\sigma$, $cm^{-1}$) |
|---|---|---|---|
| | 1625.7 | 0.56 | 19.8 |
| | 1642.1 | 0.43 | 14.2 |
| | 1660.8 | 0.84 | 16.8 |
| | 1679.1 | 0.23 | 10.5 |
| | 1692.8 | 0.13 | 12.4 |

| 3-day fibril Type-3 | Center frequency ($x_0$, $cm^{-1}$) | Amplitude (a) | Width ($\sigma$, $cm^{-1}$) |
|---|---|---|---|
| | 1623.9 | 0.46 | 19.5 |
| | 1641.8 | 0.46 | 15.0 |
| | 1661.3 | 0.86 | 16.5 |
| | 1679.5.1 | 0.26 | 11.7 |
| | 1695.1 | 0.03 | 5.6 |
| | 1736.2 | 0..24 | 16.8 |

| 5-day fibril | Center frequency ($x_0$, $cm^{-1}$) | Amplitude (a) | Width ($\sigma$, $cm^{-1}$) |
|---|---|---|---|

| | | | |
|---|---|---|---|
| | 1621.0 | 0.23 | 18.5 |
| | 1641.8 | 0.54 | 16.8 |
| | 1660.8 | 0.69 | 15.6 |
| | 1678.0 | 0.56 | 14.7 |
| | 1695.2 | 0.39 | 17.2 |
| | 1720.2 | 0.13 | 18.8 |

| 10-day fibril | Center frequency ($x_0$, $cm^{-1}$) | Amplitude (a) | Width ($\sigma$, $cm^{-1}$) |
|---|---|---|---|
| | 1620.0 | 0.38 | 15.6 |
| | 1639.8 | 0.65 | 17.1 |
| | 1661.7 | 0.82 | 17.0 |
| | 1679.2 | 0.19 | 11.0 |
| | 1692.4 | 0.18 | 17.0 |
| | 1723.6 | 0.05 | 14.2 |

| 15-day fibril | Center frequency ($x_0$, $cm^{-1}$) | Amplitude (a) | Width ($\sigma$, $cm^{-1}$) |
|---|---|---|---|
| | 1622.2 | 0.27 | 17.9 |
| | 1642.2 | 0.57 | 17.1 |
| | 1662.2 | 0.77 | 16.2 |
| | 1678.0 | 0.29 | 12.4 |
| | 1690.8 | 0.31 | 18.1 |
| | 1721.4 | 0.06 | 19.4 |
